# The HtrA chaperone monitors sortase-assembled pilus biogenesis in *Enterococcus faecalis*

**DOI:** 10.1101/2023.11.20.567783

**Authors:** Adeline M.H. Yong, Cristina Colomer Winter, Kelvin K. L. Chong, Iris Hanxing Gao, Artur Matysik, Swaine L. Chen, Kimberly A. Kline

## Abstract

Sortase-assembled pili contribute to virulence in many Gram-positive bacteria. In *Enterococcus faecalis*, the endocarditis and biofilm-associated pilus (Ebp) is polymerized on the membrane by sortase C (SrtC) and attached to the cell wall by sortase A (SrtA). In the absence of SrtA, polymerized pili remain anchored to the membrane (i.e. off-pathway). Here we show that the high temperature requirement A (HtrA) bifunctional chaperone/protease of *E. faecalis* is a quality control system that clears aberrant off-pathway pili from the cell membrane. In the absence of HtrA and SrtA, accumulation of membrane-bound pili leads to cell envelope stress and partially induces the regulon of the ceftriaxone resistance-associated CroRS two-component system, which in turn causes hyper-piliation and cell morphology alterations. Inactivation of *croR* in the Δ*srtA*Δ*htrA* background partially restores the observed defects of the Δ*srtA*Δ*htrA* strain, supporting a role for CroRS in the response to membrane perturbations. Moreover, absence of SrtA and HtrA decreases basal tolerance of *E. faecalis* against cephalosporins and daptomycin. The link between HtrA, pilus biogenesis and the CroRS two-component system provides new insights into the *E. faecalis* response to endogenous membrane perturbations.

**Author summary:** To explore the role of the HtrA chaperone/protease in *E. faecalis* off-pathway pilus clearance, we deleted *htrA* in an *E. faecalis* OG1RF Δ*srtA* strain known to retain polymerized pili on the cell membrane. Cells in the Δ*srtA*Δ*htrA* background are hyper-piliated, possess altered morphology, and are more susceptible to cell envelope-targeting antibiotics as compared to the parent OG1RF strain. RNA sequencing of the Δ*srtA*Δ*htrA* strain revealed transcriptional changes reminiscent of a membrane stress response. This response was pilus-dependent and contained several members of the CroR regulon. Inactivation of the response regulator CroR in the Δ*srtA*Δ*htrA* background restored (at least partially) piliation and cell morphology but not antibiotic susceptibility, linking CroR for the first time to pilus biogenesis and endogenous cell envelope stress.

## Introduction

Sortase-assembled pili are multi-subunit fibrillar structures that are assembled and covalently attached to the cell wall by sortase enzymes (1). Conserved among many Gram-positive bacteria, including *Enterococcus faecalis, Streptococcus pneumoniae,* and *Corynebacterium diphteriae*, sortase-assembled pili are often important virulence factors that contribute to distinct steps of the infectious process, such as host tissue adherence and biofilm formation. In *E. faecalis*, the endocarditis and biofilm-associated pilus (Ebp) operon consists of three pilin genes, *ebpA, ebpB* and *ebpC,* and a pilus-specific sortase (*srtC*) whose expression is controlled by a second promoter (2). Similar to other piliated Gram-positive bacteria, pilin monomers are translocated across the membrane via the Sec secretion machinery, assembled on the membrane into fibers by sortase enzyme C (SrtC), and subsequently attached to the cell wall by the house-keeping sortase enzyme A (SrtA), encoded elsewhere on the chromosome (3). Pili are typically expressed by only a fraction of the cell population (10-40%) (2, 4). However, the percentage increases in response to host-related environmental stimuli, including serum, bicarbonate, and fibrinogen (5-8). Ebp regulation occurs at the transcriptional level, where it is directly, positively regulated by EbpR (9). Indirectly, *ebp* transcription is activated by the RNase J2 (*rnjB*) and repressed by the quorum-sensing response regulator FsrA, in both cases through regulation of *ebpR* expression (6, 9-12).

Though a substantial number of studies have characterized pili and their contribution to infection, most studies focus on pilus regulation and biogenesis under optimal laboratory growth conditions, which may not fully recapitulate physiological conditions within the host. For example, during infection, endogenous cellular stresses or harsh exogenous conditions such as gastric acids, gut bile salts, or oxidative stress give rise to protein folding defects that can compromise the correct assembly and localization of proteins including pili, thus interfering with their function (13). This was first described in *Escherichia coli,* where misfolded pilins aggregate in the periplasm and are driven ‘off-pathway’ rather than being assembled into pili (14). Follow-up studies demonstrated that overexpression of misfolded, aggregated pilins activate two different two-component systems (TCS) in *E. coli*: Cpx and Bae (14-19). While both systems synergistically induce rapid expression of the aggregate-resolving Sly chaperone, only Cpx activates DegP, a conserved serine protease of the HtrA (<u>H</u>igh-<u>T</u>emperature <u>R</u>equirement <u>A</u>) family that degrades misfolded pilins.

It is currently unknown how Gram-positive bacteria respond to and clear aggregated pilins (20). Analogous to *E. coli*, Gram-positive bacteria also encode HtrA proteins, which primarily function as proteases involved in degradation of misfolded proteins during stress conditions (21). HtrA proteins were also shown to act as chaperones during protein assembly, and during targeting to the cell surface or the extracellular milieu. In this work, we hypothesized that the membrane-anchored serine protease HtrA might also clear off-pathway pili in Gram-positive bacteria. Using *E. faecalis* as a model organism for Gram-positive sortase-assembled pilus biogenesis, our studies confirm the role of HtrA in Ebp pili quality control. Accumulation of membrane-bound pili in the absence of HtrA perturbed the cell envelope and partially activated the CroRS two-component system, leading to hyper-piliation and changes in cell morphology. Here, we uncover a novel link between HtrA, sortase-assembled pili, and the enterococcal antibiotic-responding TCS CroRS in a Gram-positive organism.

## Results

### *E. faecalis* HtrA is not required for growth, but supports persistence in a wound infection model

Since the HtrA protein of *E. coli* was first discovered as a heat shock-inducible serine protease, and the HtrA homologues of some Gram-positive bacteria were later shown to be heat-inducible such as in *Bacillus subtilis* and *Lactobacillus helveticus* (22-24), we first determined if HtrA is involved in the heat shock response of *E. faecalis*. We generated an *E. faecalis* OG1RF Δ*htrA* deletion mutant and screened the strain for growth defects when challenged with stresses known to cause protein misfolding and aggregation (13). Incubation of WT and Δ*htrA* cells at 30°C, 37°C and 42°C did not yield significant planktonic nor biofilm growth defects compared to the WT strain (**Fig 1A & Fig S1A**). In parallel, whole lysate immunoblots of cells grown at these temperatures showed that HtrA levels only decreased at 50°C (**Fig 1B**) albeit no growth defect was observed at this temperature (**Fig S1A**), altogether suggesting that HtrA is not a major contributor to the heat shock response of *E. faecalis* under the tested laboratory growth conditions. Since other types of environmental stresses also cause protein misfolding, we challenged the *E. faecalis ΔhtrA* strain with pH, osmotic, and oxidative stress, but no growth differences were observed compared to the parent strain under laboratory growth conditions (**Fig S1B**). To investigate the strain under a more relevant host environment, we tested the Δ*htrA* mutant for fitness in a competitive mouse wound infection model available in the lab (25). Wounds were infected with a total of ∼10^6^ CFU consisting of a 1:1 mix of either OG1RF WT/OG1X or Δ*htrA/*OG1X and harvested after 8h or 72h post-infection. These two time-points were chosen to determine if HtrA might play a role in active replication (8hpi) or in long-term persistence as previously established (72hpi) (25). We chose OG1X as the comparative strain due to its close genetic kinship to OG1RF (5 different SNPs (26)) and their different antibiotic resistance profile, enabling convenient quantification on selective media (25). We recovered a median titer of ∼10^7^ CFU/ml for all strains after 8h post-infection, indicating no differences in the acute phase of infection (**Fig 1C**). However, after 72h post-infection, we recovered ∼1log less Δ*htrA* (∼1.8 x 10^4^ CFU/ml) than OG1RF (∼1.28 x 10^5^ CFU/ml). Complementation of *htrA* in *trans* (Δ*htrA* + p*htrA*) partially restored the competitive index of the Δ*htrA* strain, suggesting that HtrA might promote *E. faecalis* persistence in wounds. Overall, lack of *htrA* appeared to have negligible effects on *E. faecalis* growth *in vitro* and played a minor role in *in vivo* wound persistence.

**Fig 1.**
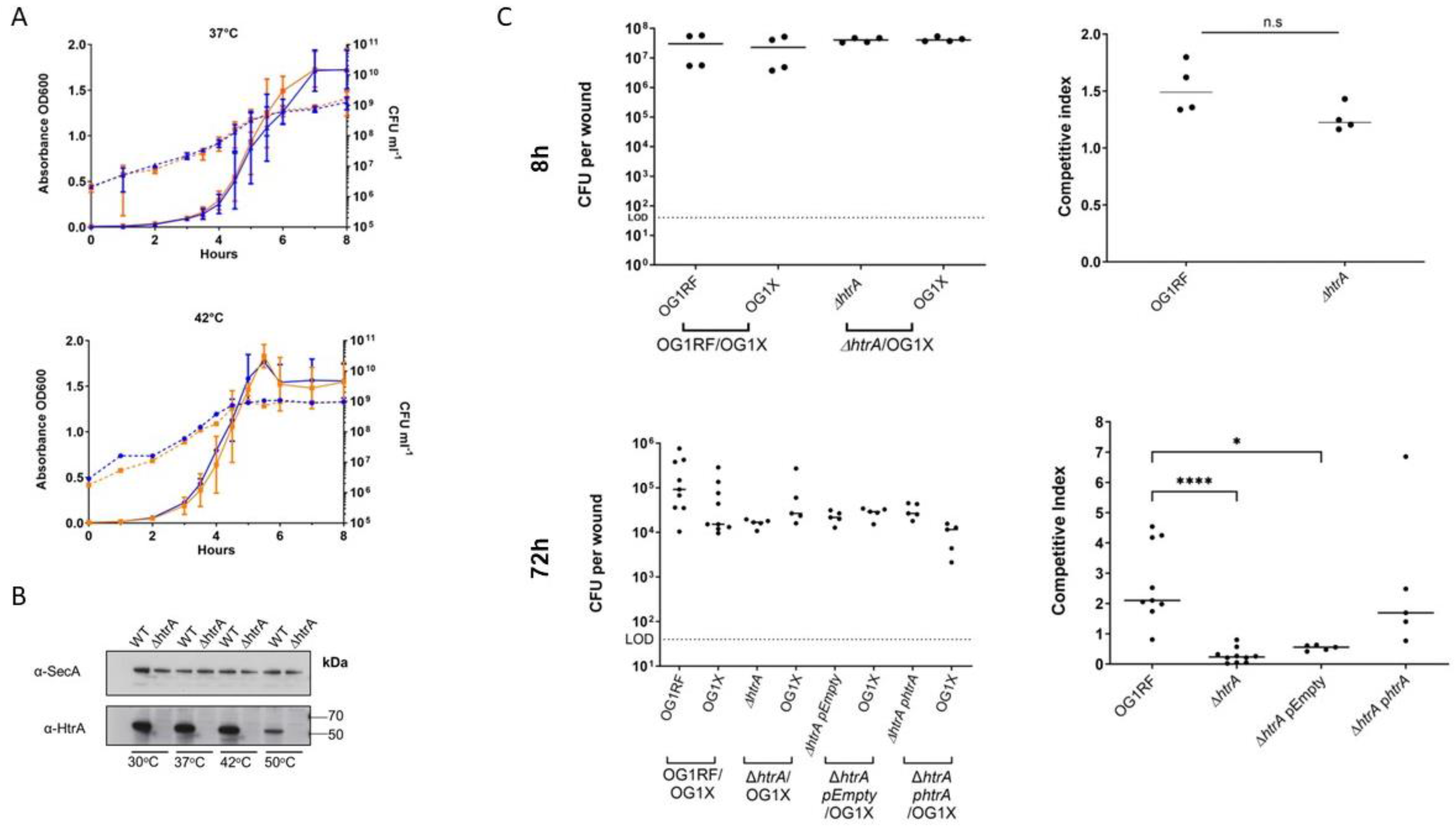
HtrA supports *E. faecalis* persistence in wounds. **(A)** Growth phenotypes of *E. faecalis* WT (orange) and Δ*htrA* (blue). Growth curves performed in BHI broth at 37°C and 42°C. Representative data of three independent experiments. CFU counts (CFU ml^-1^) are represented as dashed lines; OD_600_ readings are represented as solid lines. Standard deviation is indicated by bars. **(B)** Western blots of whole cell lysates of OG1RF WT and Δ*htrA* grown at different temperatures. To detect HtrA, affinity purified α-HtrA was used. α-SecA was used as a loading control. **(C)** Wounds were infected with a 1:1 ratio of *E. faecalis* strains OG1X/OG1RF WT or OG1X/OG1RF Δ*htrA*, at 10^6^ CFU per inoculum, and harvested at 8 hpi or 72 hpi. Recovered bacteria were enumerated on selective media for each strain. Brackets represent strain pairs that were co-infected. Each dot represents a mouse. Competitive index was calculated using the final CFU ratio of OG1X with OG1RF WT or Δ*htrA* (output) over the initial CFU ratio of OG1X with OG1RF or Δ*htrA* (input). Solid horizontal line indicates the median. The limit of detection (LOD) of 40 CFU is indicated. For 8h N=1, n=4. For 72h N=1, n=10 for OG1RF and Δ*htrA,* n=5 for Δ*htrA* p*Empty* and Δ*htrA*p*htrA.* Statistical analysis was performed using the Kruskal-Wallis test with Dunn’s post-test to correct for multiple comparisons. **** P≤0.0001, * P<0.05.

### The HtrA chaperone contributes to removal of off-pathway Ebp pili

Since the main HtrA homolog in *E. coli*, DegP, has been shown to remove misfolded pili in *E. coli* (14), we asked whether HtrA might be involved in pili biogenesis in enterococci. In *E. faecalis*, the housekeeping sortase A (SrtA) efficiently anchors polymerized pili on the cell wall (27). When *srtA* is inactivated (Δ*srtA*), polymerized pili remain anchored on the cell membrane and protrude through the cell wall (27, 28). We hypothesized that HtrA might be involved in clearing aberrant membrane-anchored pili in *E. faecalis*. Thus, we predicted that removal of *htrA* in the Δ*srtA* background (Δ*srtA*Δ*htrA*) would lead to accumulation of pili on the cell membrane. To test this, we performed immunoblot analysis on cell wall and protoplast fractions of wild type (WT), Δ*srtA, ΔhtrA*, and Δ*srtA*Δ*htrA* strains with anti-EbpC immune serum. As expected for sortase-dependent cell wall-anchored pili (27), the major pilin subunit EbpC was found in the cell wall fraction of the WT strain (**Fig S2A**) and absent in the protoplast fraction (**Fig 2AB**). In line with previous results, EbpC was absent from the cell wall fraction of *srtA* mutants (**Fig S2A & S2D**) and remained instead on the protoplast fraction (**Fig 2AB**). Single deletion of *htrA* did not alter EbpC cell fraction localization as the pilus subunit was found in the cell wall fraction similar to the WT strain. Interestingly, we found that EbpC levels were 4.6-fold higher in the protoplast fraction of the Δ*srtA*Δ*htrA* mutant strain than in the Δ*srtA* strain, whereas expression of a control protein (SecA) was unaffected (**Fig 2A, B**). We observed similar trends with minor subunits EbpA and EbpB (**Fig S2B. C, E, F**), and complementation of either *srtA* or *htrA* in the Δ*srtA*Δ*htrA* background restored, at least in part, pilus levels and cell fraction location (**Fig 2A, B**).

**Fig 2.**
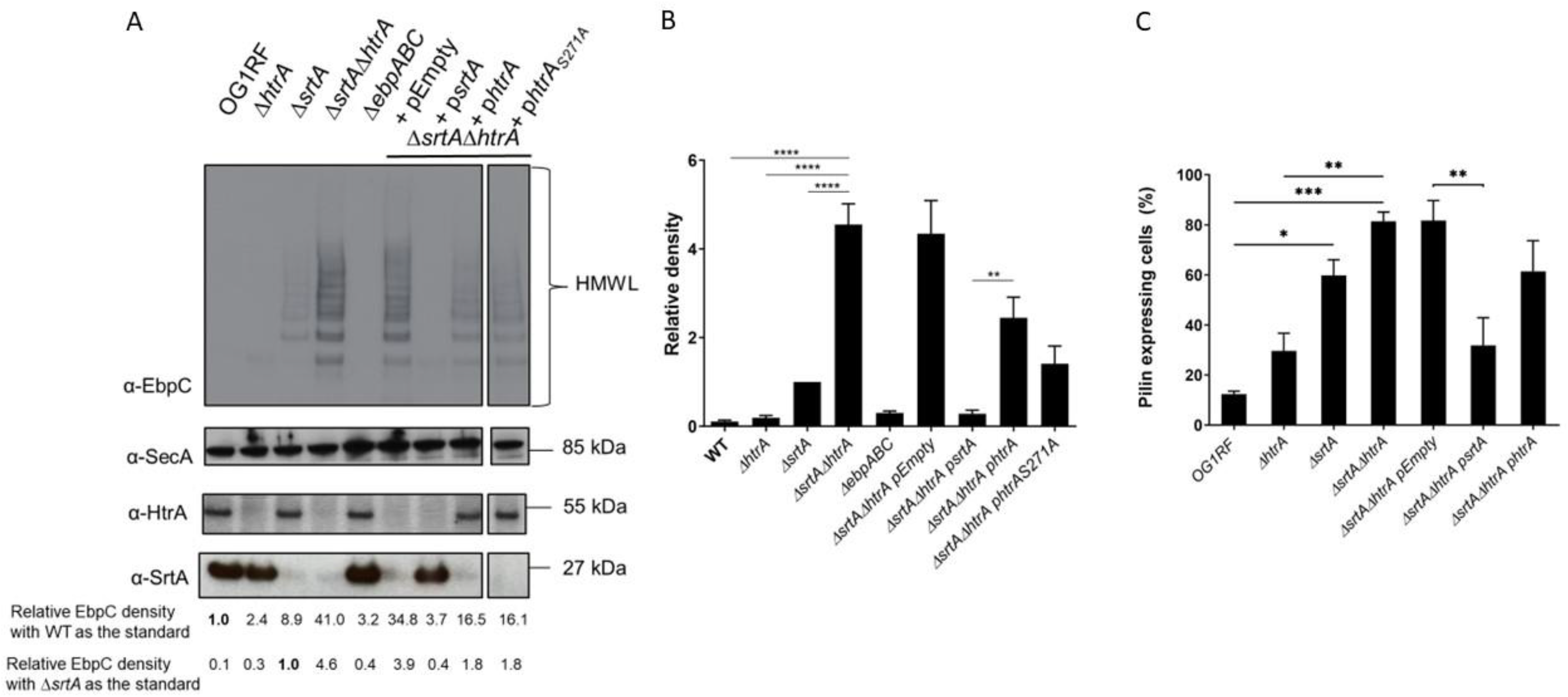
Loss of *htrA* increases EbpC protein levels in the protoplast fraction. **(A)** Immunoblot was performed with α-EbpC on protoplast fractions of WT, Δ*htrA, ΔsrtA, ΔsrtAΔhtrA, ΔebpABC* strains, as well as Δ*srtA*Δ*htrA* carrying p*Empty* (vector control), p*srtA,* p*htrA* or p*htrA_S271A_*. Top blot shows typical pilus high molecular weight ladders (HMWL) and bottom blots show loading and strain controls, respectively, using α-SecA, α-HtrA and α-SrtA. Relative EbpC densities differences were calculated with WT or Δ*srtA* EbpC expression as the standard. **(B)** Statistical analyses of relative EbpC density from four independent immunoblots, using Δ*srtA* as the comparison standard, are represented as bar graphs with the standard error of the mean. ** P ≤ 0.01; **** P ≤ 0.0001; P ≥ 0.05 differences not significant. (**C**) Statistical analyses of pilus-expressing cells of WT, Δ*htrA, ΔsrtA, ΔsrtAΔhtrA*; Δ*srtA*Δ*htrA* strains carrying p*Empty* (vector control), p*srtA* or p*htrA*, labeled with α-EbpC immune serum and Alexa Fluor® 568 secondary antibody. Mean results are represented as bar graphs with standard error of mean. * P < 0.05; ** P ≤ 0.01; *** P≤0.001, **** P ≤ 0.0001. Combined data from three independent experiments were shown.

HtrA enzymes often display dual chaperone and protease functions (29, 30). Since Ebp remained bound to the protoplast fraction of the single Δ*srtA* and the double Δ*srtA*Δ*htrA* strains, but total EbpC levels were significantly higher upon *htrA* inactivation (**Fig 2**), we hypothesized that the HtrA serine protease activity may directly degrade membrane-bound Ebp. We designed an HtrA expression plasmid with a single amino-acid change in the conserved serine (S271A) of the proteolytic active site of the enzyme (p*htrA*_S271A_), inactivating the protease function without altering the chaperone function (21, 31). Immunoblot analysis revealed a decrease in EbpC levels in the Δ*srtA*Δ*htrA* strain complemented with p*htrA*_S271A_, similar to Δ*srtA*Δ*htrA* complemented with the wild-type p*htrA* (**Fig 2AB**), indicating that the HtrA protease activity is dispensable and its chaperone activity is sufficient to remove the accumulated pili from the cell membrane.

Approximately 10-40% of WT cells grown under laboratory conditions in TSBG (10-20%) or BHI (20-40%) media express Ebp (2, 4). Though EbpR-dependent regulation of *E. faecalis* pili is well characterized, it is currently unknown why only a subpopulation of cells express pili. Inactivation of *srtA* was previously shown to correlate with increased total pilus abundance (27). However, it was unknown whether the increased Ebp levels observed on the immunoblot of Δ*srtA*Δ*htrA* cells reflected the frequency of piliated cells in the total cell population, or hyper-piliation on a single cell level. Immunofluorescence of EbpC labeled cells grown in TSBG revealed that 80% of Δ*srtA*Δ*htrA* cells in the population expressed pili as compared to WT (15%), while Δ*srtA* (60%) and Δ*htrA* (30%) displayed incremental increases in total population piliation (**Fig 2C**). Complemented strains reverted to single mutant piliation levels. These results indicate that efficient pilus cell wall-anchoring by SrtA and monitoring by HtrA is an important aspect in the regulation of cell piliation in a given population.

### Accumulation of membrane-bound pili elicit broad transcriptional changes

To investigate the transcriptional response to accumulation of membrane-bound pili, we performed RNA sequencing of OG1RF WT, Δ*srtA*, Δ*htrA* and Δ*srtA*Δ*htrA*. When compared to OG1RF WT, single inactivation of *htrA* led to differential expression of 4 genes (3 hypothetical genes and flavocytochrome C), while deletion of *srtA* altered transcription of 11 genes, including repression of PTS sugar transport genes, and up-regulation of arginine metabolism and the quorum sensing transcriptional regulator *fsrA* (**Table S4**). Strikingly, simultaneous deletion of *srtA* and *htrA* changed the expression of 307 genes (106 up-regulated, 201 down-regulated).

Accumulation of misfolded, aggregated pili has been shown to perturb the inner cell membrane of *E. coli* (14-16). In *E. coli*, inner membrane perturbations are sensed by the Cpx two <u>c</u>omponent <u>s</u>ystem (TCS), which orchestrates a stress response to return to membrane homeostasis. The Cpx regulon includes repression of non-essential lipoproteins (such as solute transport systems), and concomitant activation of protein folding enzymes (such as DegP and Spy) and peptidoglycan-modifying enzymes (15, 32). We hypothesized that accumulation of membrane-bound pili could similarly perturb the cell membrane of *E. faecalis.* We thus searched for commonalities between the *E. coli* Cpx regulon and the *E. faecalis* transcriptional response to aggregated, accumulated pilins. To streamline our efforts, pairwise transcriptome comparisons between the double mutant strain and either the WT or the single *htrA* or *srtA* mutant strains identified 164 genes that were differentially regulated in all three comparisons (p<0.05; FDR<0.05), suggesting that these genes may be specifically and consistently involved in the response to accumulation of membrane-bound pili (**Fig 3A, Table S5**). We classified these 164 consensus genes into predicted functional groups for further analysis. The log_2_ fold change in mRNA levels in Δ*srtA*Δ*htrA* are displayed relative to Δ*srtA* as an example and are shown as a dot plot (**Fig 3B**).

**Fig 3.**
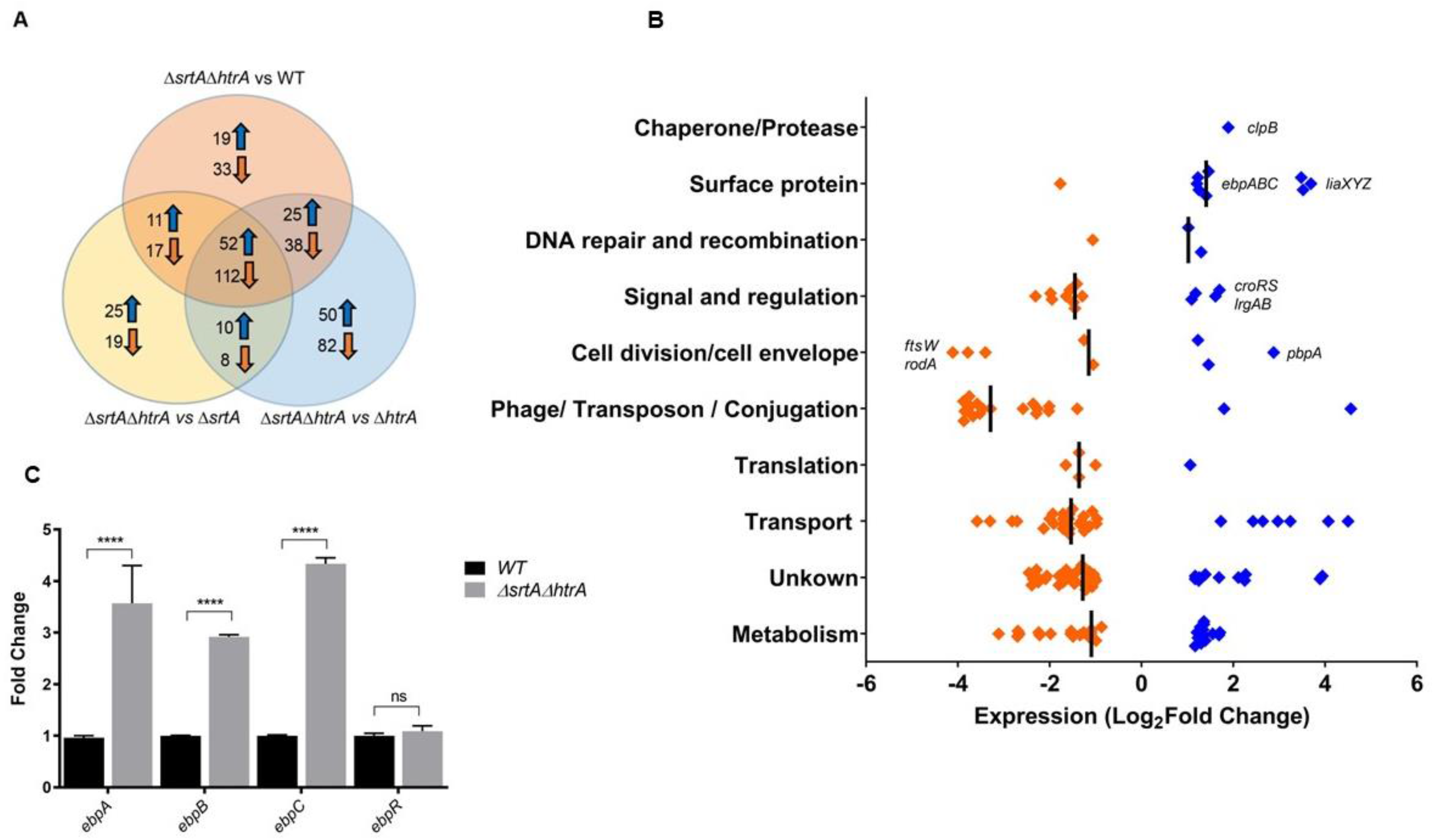
Global transcriptional changes due to absence of *htrA* and *srtA*. **(A)** Venn diagram showing a total of 164 genes found to be differentially regulated specifically during pili accumulation on the cell membrane (i.e. in the absence of both *srtA* and *htrA*). Genes that are only found in one or two of the indicated groups are in the non-overlapping regions. The total number of differentially expressed genes in each comparison is indicated inside each group. **(B)** The log_2_ fold change in mRNA levels of the core 164 genes that are differentially expressed in the Δ*srtA*Δ*htrA* strain. Values displayed as a dot plot correspond to differences between Δ*srtA*Δ*htrA* and Δ*srtA*. Significant genes were determined by the Bioconductor package EdgeR (P < 0.05; FDR < 0.05). Genes with negative log_2_ fold change are colored orange; genes with positive log_2_ fold change are colored blue. Genes of interest were labeled for easier identification. **(C)** qRT-PCR analysis of *ebpABCR* expression in the WT (black bars) and Δ*srtA*Δ*htrA* (grey bars). qRT-PCR was performed in biological triplicates and analyzed by the ΔΔ_CT_ method, using *gyrB* as a housekeeping gene. Fold change indicates the change in *ebpA, ebpB*, *ebpC* and *ebpR* transcription compared to WT. Statistical analysis was performed by 2-way ANOVA and Turkey’s multiple comparison tests using GraphPad. **** P ≤ 0.0001, P ≥ 0.05 differences not significant (ns).

In line with the *E. coli* Cpx inner membrane stress response (15, 33, 34), a significant number of repressed genes clustered in the transport and metabolism categories, including non-essential sugar and amino acid transporters. Moreover, genes involved in peptidoglycan turnover were induced, including the penicillin-binding protein *pbpA* (∼1.7 log_2_-fold), the murein hydrolase regulators *lrgAB,* and the *croRS* TCS (∼ 1.5 log_2_-fold). <u>Cr</u>oRS [short for “ceftriaxone resistance”] typically regulates penicillin-binding proteins in response to antibiotic-induced cell wall damage (35-37). In the protease/chaperone category we only observed upregulation of *clpB* (∼1.7 log_2_-fold), a key chaperone that removes protein aggregates in several bacterial pathogens (13, 38). In contrast to *E. coli*, differentially expressed genes that clustered in the translation category were generally repressed. Overall, these results suggested that off-pathway pili also cause cell envelope perturbations in *E. faecalis*, but that the coping strategies utilized by this Gram-positive bacterium partially differ from *E. coli*. Transcriptional analysis of the double Δ*srtA*Δ*htrA* strain harbored additional findings supporting membrane stress. For example, accumulation of off-pathway pili led to strong upregulation of the *liaXYZ* operon (∼3 log_2_-fold) involved in modulation of the membrane stress-sensing TCS LiaFSR. Differential expression of cell division genes such as putative *ftsW* and *rodA* was also detected. Consistent with the increased pilus expression observed in this strain, transcription of the pilin genes *ebpA, ebpB* and *ebpC* was significantly induced (∼1.6 log_2_-fold RNA-Seq, ∼3.6-fold qRT-PCR) in the Δ*srtA*Δ*htrA* mutant (**Fig 3BC**). Transcript levels of the positive regulator *ebpR,* as well as of *ebpR*-regulating *rnjB* and *fsrA,* were unchanged in the Δ*srtA*Δ*htrA* strain (**Table S4**), suggesting that induction of the *ebp* locus may be regulated independently of EbpR.

### Accumulation of off-pathway pili partially mobilizes the CroR regulon

Since TCS such as Cpx and Bae sense pilus aggregates in *E. coli* (14, 16-19), and transcription of the *croRS* TCS was induced (∼1.4 log_2_-fold RNA-Seq, ∼2.5-fold qRT-PCR) in the *E. faecalis ΔsrtAΔhtrA* strain (**Fig 3B & Fig 4A**), we sought to determine if CroRS regulated at least part of the pilus-responsive stress response. We constructed a *croR::tnΔsrtAΔhtrA* triple mutant strain and performed immunofluorescence pili staining as part of the initial characterization. Unexpectedly, *croR* inactivation significantly decreased piliation levels in the cell population (*croR::tnΔsrtAΔhtrA* (36%)) as compared to Δ*srtA*Δ*htrA* (80%) (**Fig 4B**). In line with pilus surface expression, inactivation of *croR* also decreased *ebp* transcription levels as quantified per qRT-PCR (**Fig 4C**). Complementation of *croR::tnΔsrtAΔhtrA* with *croRS* on an expression plasmid (*pcroRS-2xHA*) caused again hyper-piliation (**Fig.S3**).

**Fig 4.**
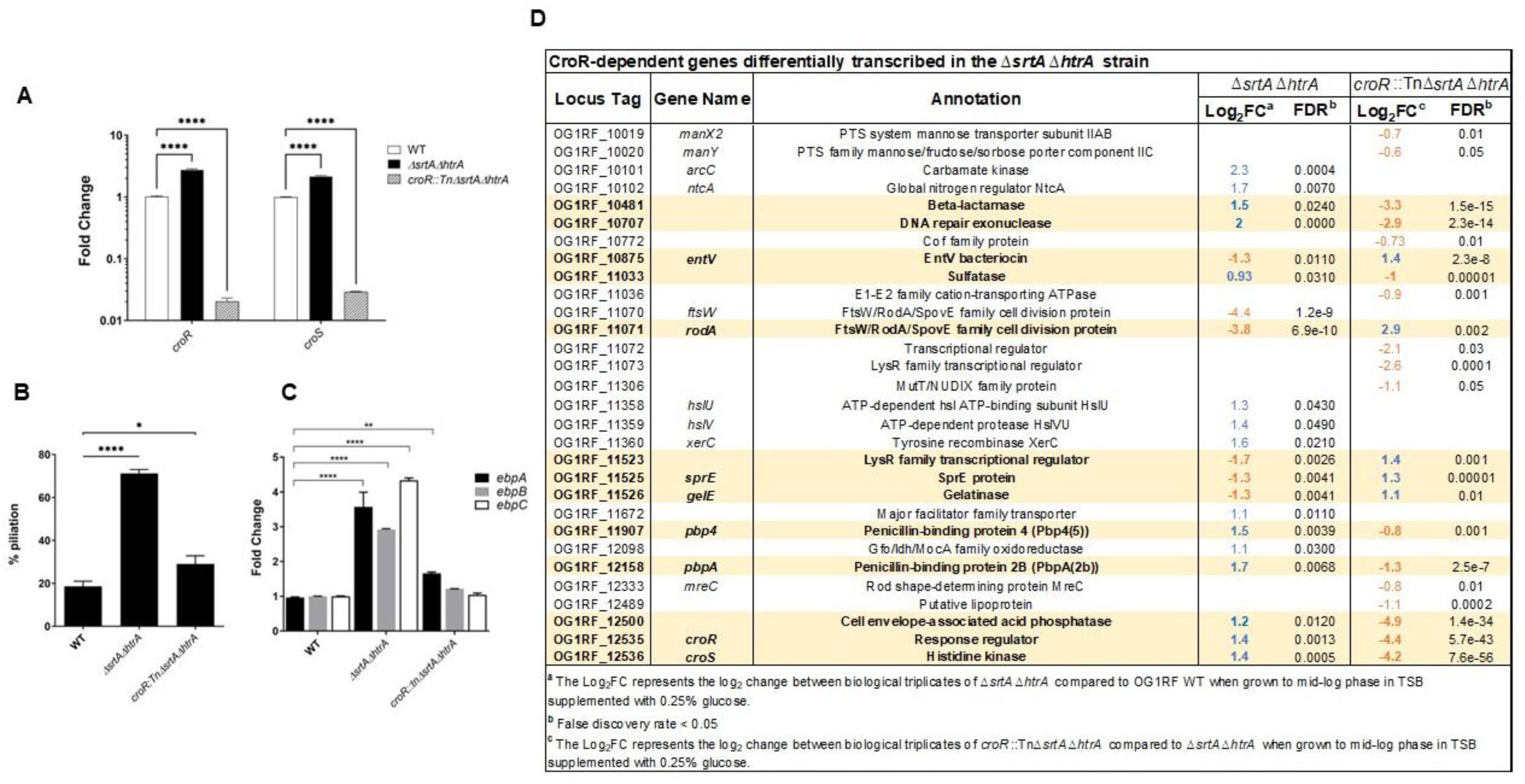
Accumulation of off-pathway pili on the cell membrane activate the CroRS system. (**A**) qRT-PCR analysis of *croRS* expression in the WT (black bars), Δ*srtA*Δ*htrA* (light grey bars), and *croR::tnΔsrtAΔhtrA* (dark grey bars). qRT-PCR was performed in biological triplicates and analyzed by the ΔΔ_CT_ method, using *gyrB* as a housekeeping gene. Fold change indicates the change in *croR* and *croS* transcription compared to WT. c*roR::tnΔsrtAΔhtrA* was used as the negative control for *croRS* transcription. Statistical analysis was performed by the 1-way ANOVA and Tukey’s comparison test using GraphPad. **** P ≤ 0.0001. (**B**) Statistical analysis of pili expressing cells labeled with α-EbpC immune serum and Alexa Fluor® 568 secondary antibody. Results are represented as bar graphs with standard error of mean. Combined data from three independent experiments were shown. Statistical analysis was performed by the 2-way ANOVA and Turkey multiple comparison test using GraphPad. * P ≤ 0.05; *** P ≤ 0.001; **** P ≤ 0.0001; P > 0.05, differences not significant (ns). (**C**) qRT-PCR analysis of *ebpA* (black bars), *ebpB* (light gray bars) and *ebpC* (white bars) expression in the WT, Δ*srtA*Δ*htrA* and *croR::tnΔsrtAΔhtrA* strains. qRT-PCR was performed in biological triplicates and analyzed by the ΔΔ_CT_ -method using *gyrB* as a housekeeping gene. Fold change indicates the change in *ebpABC* gene transcription compared to WT. Statistical analysis was performed by the 2-way ANOVA and Turkey multiple comparisons test using GraphPad. **** P ≤ 0.0001. (**D**) List of CroR-dependent genes identified by Timmler *et al* in bacitracin-treated OG1 (35) that are differentially expressed in the Δ*srtA*Δ*htrA* strain (compared to OG1RF WT), in the *croR::tnΔsrtAΔhtrA* strain (compared to Δ*srtA*Δ*htrA),* or in both. Log_2_FC values are indicated in blue for upregulated genes and orange for downregulated genes. Genes highlighted in bold font and yellow background are restored, at least in part, to WT levels upon *croR* inactivation.

Basal transcription of TCS genes under unstressed conditions is typically low. When activated (i.e. usually phosphorylated) by a given signal, TCS response regulators can activate their own transcription in a positive feedback loop until the stress is resolved (39). Upregulation of *croRS* in the Δ*srtA*Δ*htrA* strain suggests that membrane overloading with off-pathway pili might activate the CroRS system. If true, we would expect to see at least part of the CroR regulon embedded in the transcriptome of the Δ*srtA*Δ*htrA* mutant, and inactivation of CroR would restore CroR-dependent genes to WT levels. We performed RNA-Seq of the *croR::tnΔsrtAΔhtrA* triple strain and compared it to the transcriptome of the Δ*srtA*Δ*htrA* double mutant (**Table S6, Table S4**), while also scanning for the signature of the CroR regulon in the Δ*srtA*Δ*htrA* strain. Of note, to avoid strain specific differences we used the CroR regulon identified using bacitracin-treated OG1 as the background strain as our reference (35). Our analysis uncovered that ∼24% of CroR-dependent genes (21 out of 88 genes) were differentially expressed in the Δ*srtA*Δ*htrA* strain (**Fig 4D**). Of these, ∼52% were restored, at least in part, to wild-type levels upon *croR* inactivation, including hallmark CroR-dependent genes such as the cephalosporin low affinity penicillin-binding proteins *pbp4* and *pbpA* (35).

### Aberrant cell morphology of the Δ*srtA*Δ*htrA* strain is Ebp- and CroR-dependent

Transcriptomic analysis revealed several putative cell division genes that were differentially expressed in the Δ*srtA*Δ*htrA* strain (**Fig 3B**) and that were partially restored in the triple *croR::tnΔsrtAΔhtrA* strain (**Fig 4D**). To determine the presence of cell morphology defects, we first performed phase contrast microscopy of the OG1RF parental strain and the panel of single *htrA*, *srtA* and double mutant strains. A distinct phenotype was observed in a subset of the Δ*srtA*Δ*htrA* population, characterized by the presence of chains containing 4-8 joined cells (**Fig 5A & Fig S4A**). The chaining phenotype of Δ*srtA*Δ*htrA* could be complemented with p*htrA* or p*srtA*, resulting in reversion to a diplo-ovococcal shape similar to WT (**Fig 5A**). To gain more insight into the morphology defect, which was suggestive of a defect in cell division or cell elongation at the peptidoglycan (PG) level, we visualized PG using a vancomycin-fluorescein (Van-FL) dye. Of note, vancomycin binds to the terminal D-Ala-D-Ala found on uncrosslinked PG precursors of the entire cell wall, allowing for visualization of peripheral cell wall as well as the septum (40). In addition to chaining, individual Δ*srtA*Δ*htrA* cells appeared shorter and wider than WT (**Fig 5B**), suggesting a defect in cell elongation. In addition, Van-FL labeling revealed aberrant placement of septa across 30% of the Δ*srtA*Δ*htrA* cell population (**Fig 5B & Fig S4B**), indicating that cell division was also affected. Further transmission electron microscopy confirmed that some Δ*srtA*Δ*htrA* cells had severely deformed septa (**Fig 5B & Fig S4C**). Moreover, these cells lost the ovococcal WT shape and exhibited a more spherical morphology. To quantify the cell shape, we measured the cell width of each strain and plotted it as a distribution curve. The majority of Δ*srtA*Δ*htrA* cells had a cell width of ∼0.8 µm, while WT, Δ*srtA* or Δ*htrA* cells neared 0.7 µm (**Fig 5C**). Altogether, it appeared that the simultaneous absence of *srtA* and *htrA* hampered the ability of the cell to elongate and divide, reminiscent of the LiaFSR-dependent cell membrane stress response in enterococci (41, 42).

**Fig 5.**
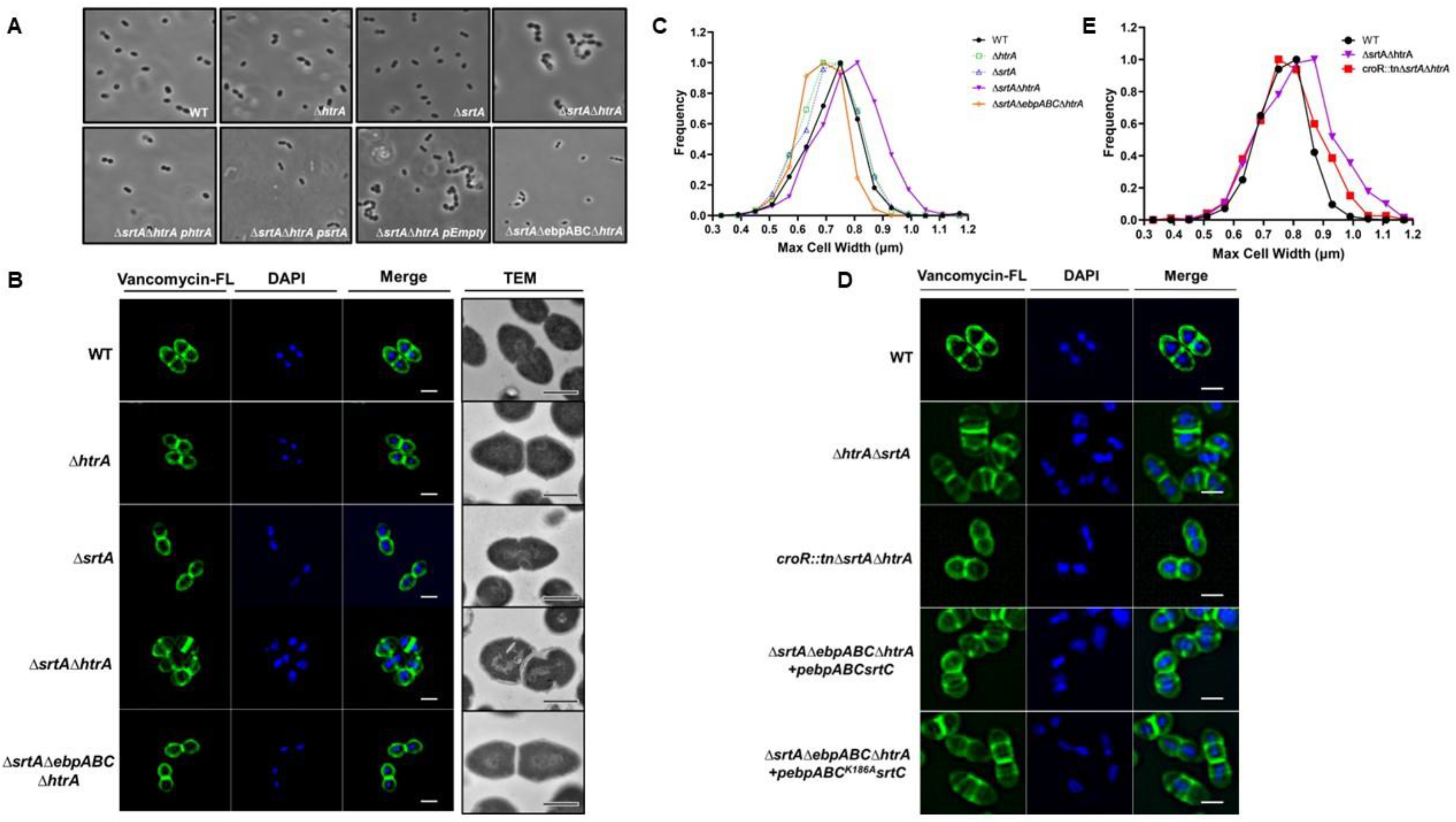
Aberrant cell morphology of the Δ*srtA*Δ*htrA* mutant is pilus- and CroR-dependent. (**A**) The morphology of the indicated *E. faecalis* strains was visualized at a magnification of 100X with a phase contrast microscope. Representative images are shown. (**B, D**) The cell wall of log-phase growing cells was stained with Vancomycin-FL conjugate (green). DAPI (blue) was used to visualize DNA. Representative images are shown. Bacterial ultrastructures were observed under TEM. Scale bars for immunofluorescence microscopy and transmission electron microscopy represent 1 µm and 500 nm, respectively. (**C, E**) The mid-cell width for each strain was determined and quantified using the MicrobeJ plugin in ImageJ and the distribution of the population is represented by a Gaussian distribution. The data was plotted using Microsoft Excel. Cells that were not in-phase were excluded from the analysis. A total of at least 200 cells was analyzed per strain.

Considering that *croR* inactivation partially restored transcription of the dysregulated cell division genes of the Δ*srtA*Δ*htrA* strain (**Fig 4D**), we hypothesized that inactivation of *croR* in the triple *croR::tnΔsrtAΔhtrA* strain would restore the division defects of the Δ*srtA*Δ*htrA*. By measuring cell width and performing cell wall staining, we observed that *croR::tnΔsrtAΔhtrA* indeed no longer exhibited morphology defects (**Fig 5DE**). Since membrane overloading with pili perturbs the cell envelope and likely triggers activation of the CroR regulon, we deleted the entire *ebp* locus in the Δ*srtA*Δ*htrA* background to remove the triggering stress source (quintuple Δ*srtA*Δ*ebpABCΔhtrA* mutant). Identical to *croR* inactivation, deletion of *ebpABC* restored cell morphology to that of the WT strain (**Figs 5AC**). Complementation of *ebpABC* in the quintuple strain triggered once again morphology defects (**Fig 5D**).

While the exact signal sensed by CroRS remains elusive, the trigger is suggested to be related to cell wall damage (35, 36, 43). Membrane-bound pili protrude through the cell wall (28) and could potentially perturb the peptidoglycan layer, activating the CroRS system. To begin to interrogate if CroRS activation was due to membrane protein overloading or peptidoglycan perturbation, we next investigated whether accumulation of monomeric pilin subunits equally induce morphological defects. We used a plasmid with a single amino-acid change in the EbpC pilin-like motif (K186A) that is necessary for pilus fiber polymerization, resulting in the expression of EbpC monomers that cannot be polymerized into full length pili (27). Complementation of Δ*srtA*Δ*ebpABCΔhtrA* with genes encoding either monomeric pilin subunits (p*ebpABC_K186A_srtC*) or polymerized pili (p*ebpABCsrtC*) both gave rise to morphology defects (**Fig 5D & S4D**), indicating that membrane perturbation was likely responsible for CroR activation.

### CroR partially induces cell morphology defects through regulation of RodA

The Δ*srtA*Δ*htrA* mutant differentially expressed a small group of genes that clustered in the cell division and cell envelope category, including *ftsH*, OG1RF_11070, OG1RF_11071, *pbp4*, *pbpA,* and *pbp2A* (**Table S4**). Transcription of *pbp4*, *pbpA*, and OG1RF_11071 were restored upon CroR inactivation (**Fig 4D**). We hypothesized that CroR might alter cell morphology defects via control of at least one of these genes. While Pbp4 and PbpA are penicillin-binding proteins linked to cell wall homeostasis, the function of OG1RF_11071 was previously unknown. OG1RF_11071 and its neighboring gene OG1RF_11070 were both annotated as FtsW/RodA/SpovE family cell division proteins. Based on BLASTp searches, the product of OG1RF_11071 shared ∼41% identity with the protein sequence of *rodA* in *Streptococcus oralis,* and OG1RF_11070 shared ∼36% identity with *ftsW* in *Streptococcus agalactiae*. We will refer to them as *ftsW* and *rodA* from here on for brevity. FtsW is a universally conserved peptidoglycan polymerase essential for septal cell wall assembly (44), while RodA is a highly conserved glycosyltransferase involved in cell wall morphogenesis (45). Both enzymes interact with penicillin-binding proteins to control cell shape and septation in other bacteria (44-47), and are predicted to be co-transcribed in a single operon in *E. faecalis*. We designed an expression plasmid containing both genes (*pftsW/rodA*) and expressed it in *trans* in the Δ*srtA*Δ*htrA* strain. The morphological defect (as measured by multiple septa) was partially alleviated in Δ*srtA*Δ*htrA pftsW/rodA*, with 9.5 ± 3% of cells still exhibiting the morphology defect, as compared to 19.3 ± 0.7 % in the double Δ*srtA*Δ*htrA* strain (**Fig 6AB**). These results suggest that CroR partially induces morphology changes through regulation of RodA, but that the morphology defect is likely multi-factorial and might involve control of partner penicillin-binding proteins such as Pbp4 and PbpA (47).

**Fig 6.**
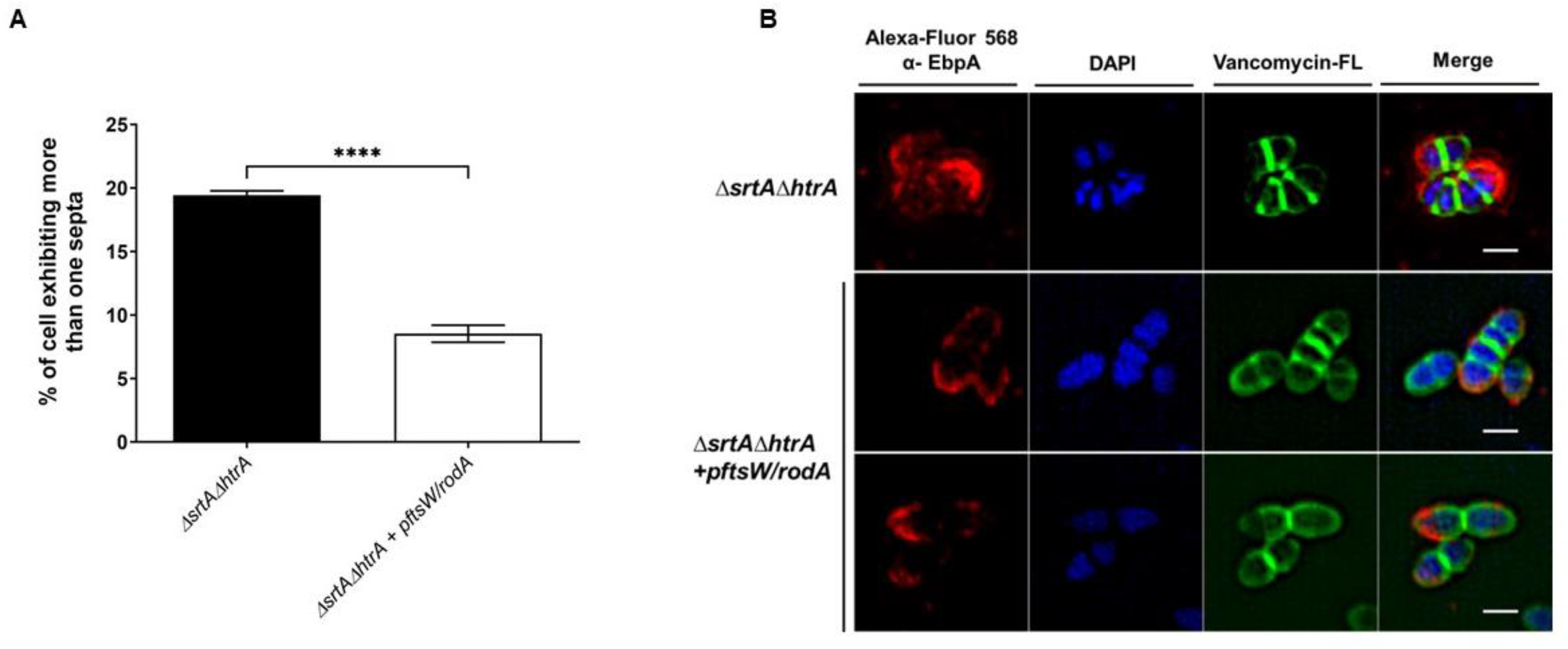
CroR partially induces cell morphology defects through transcriptional regulation of the glycosyltransferase *rodA*. **(A)** The percentage of cells exhibiting more than 1 septum per cell are represented as bars. The data was plotted using GraphPad Prism. A total of 500 cells were analyzed per strain. Statistical analysis was performed by the unpaired parametric T-test using GraphPad. **** P ≤ 0.0001. (**B**) The cell wall of Δ*srtA*Δ*htrA* and Δ*srtA*Δ*htrA* + p*ftsW/rodA* was stained with Van-FL conjugate (green), blocked and incubated with α-EbpC immune serum coupled to Alexa Fluor® 568 antibody (red). DAPI (blue) was used to visualize DNA. Representative images were shown to illustrate the partial morphology restoration. Scale bars represent 1 µm.

### Presence of the non-cognate CisS histidine kinase in the absence of CroS sustains cell morphology defects

Since CroR can be phosphorylated by the non-cognate histidine kinase CisS in the absence of CroS (43), we asked if CroR-CisS crosstalk was also present in OG1RF during endogenous membrane stress. First, we constructed a triple *croS::tnΔsrtAΔhtrA* strain and examined cells for restoration of piliation and morphology defects as a readout for CroR activation. Inactivation of the cognate *croS* histidine kinase only partially restored piliation levels and did not revert morphology defects (**Fig 7ABC**), suggesting that CroR may still be activated by CisS, hence resulting in morphological defects. To further address this hypothesis, we created a quadruple Δ*cisSΔcroSΔsrtAΔhtrA* deletion mutant to remove both sensor histine kinases and quantified the cell width of the panel of strains. The Δ*srtA*Δ*htrA* and *croS::tnΔsrtAΔhtrA* strains were on average wider (∼1.07 µm and ∼1.12 µm) than the WT and *croR::tnΔsrtAΔhtrA* strains (∼0.97 µm and ∼1.0 µm). Double inactivation of the two histidine kinases in the Δ*cisSΔcroSΔsrtAΔhtrA* strain restored average cell width to ∼0.94 µm (**Fig 7C**), suggesting the existence of CroR-CisS crosstalk under these conditions.

**Fig 7.**
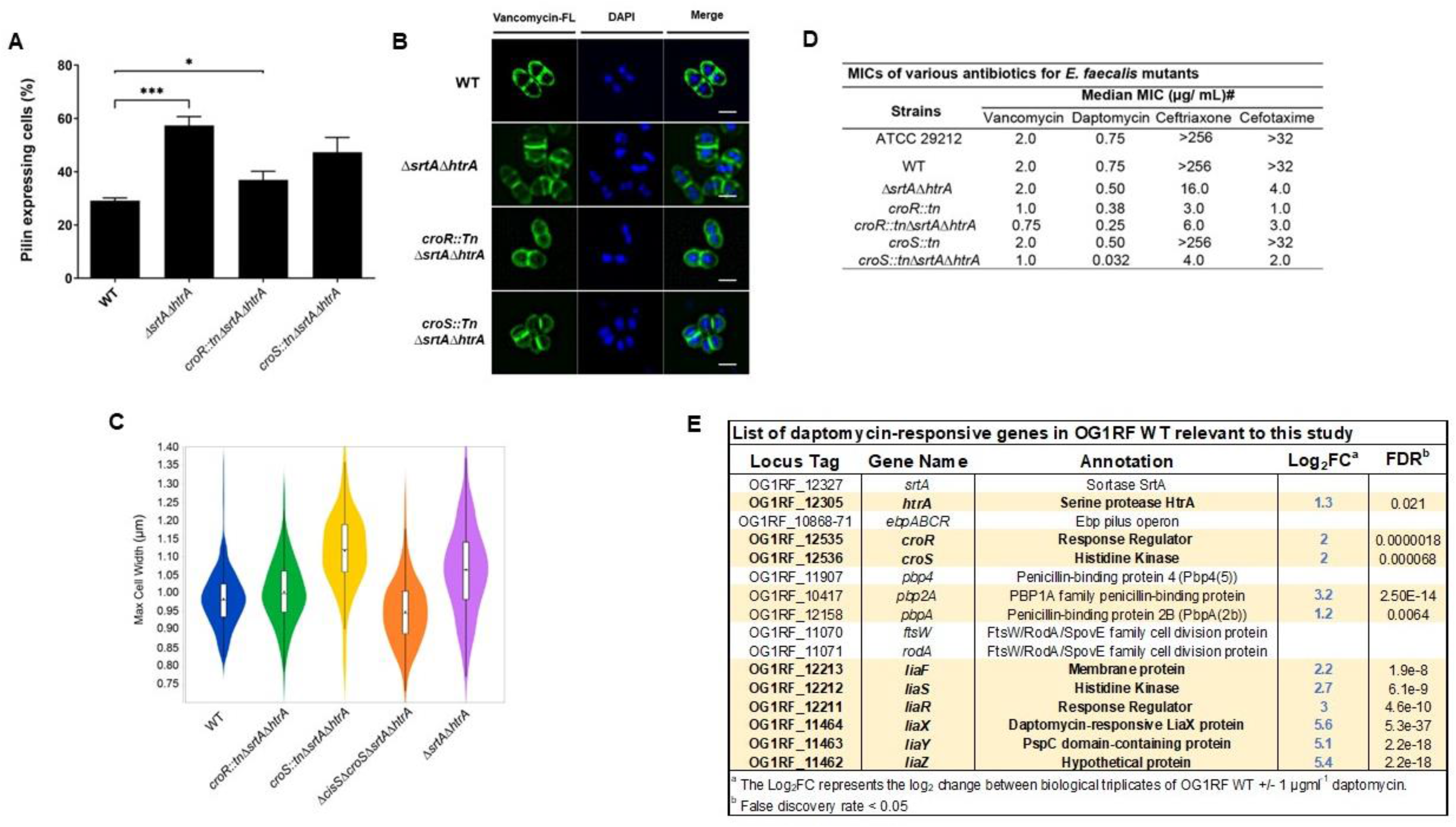
CisS support cell morphology defects and lack of *srtA* and *htrA* leads to antibiotic susceptibility. **(A)** Statistical analysis of pili expressing cells labeled with α-EbpC immune serum and Alexa Fluor® 568 secondary antibody. Results are represented as bar graphs with standard error of mean. Combined data from three independent experiments were shown. Statistical analysis was performed by the 2-way ANOVA and Turkey multiple comparison test using GraphPad. *** P ≤ 0.001; **** P ≤ 0.0001; P > 0.05, differences not significant (ns). **(B)** The cell wall was stained with Vancomycin-FL conjugate (green). DAPI (blue) was used to visualize DNA. Representative images were shown. Scale bars represent 1 µm **(C)** The maximum cell width of the various mutant strains was measured and quantified using MicrobeJ plugin in ImageJ and are represented as violin plot. Cells that were not in-phase were excluded from the analyses. A total of at least 500 cells were sampled per strain. Dot represents the mean; horizontal line represents the median; box represents the interquartile region **(D)** Minimal inhibitory concentration of various antibiotics for *E. faecalis* mutants. ^#^ indicates the median MICs reported from 2 biological replicates. ATCC 29212 is the positive control strain that fits the standard MIC of gentamicin (4-16 µg/mL) **(E)** List of selected genes relevant to this study and their log_2_-fold change in transcription in OG1RF WT treated with subinhibitory concentrations of daptomycin. Differentially expressed genes are highlighted in bold font and yellow background.

### Absence of *srtA* and *htrA* leads to cephalosporin and daptomycin antibiotic susceptibility

*E. faecalis* is a leading cause of antibiotic-resistant infections, partially due to an intrinsic tolerance to cell wall-active antibiotics (48). One of the systems used by *E. faecalis* to sustain high basal tolerance to β-lactams is the CroRS TCS, in which phosphorylated CroR upregulates cephalosporin tolerant penicillin-binding proteins such as Pbp4 and PbpA in response to antibiotic treatment (35, 36, 43, 49). Since aberrant accumulation of Ebp pili triggered activation of CroRS in the Δ*srtA*Δ*htrA* strain, we hypothesized that the Δ*srtA*Δ*htrA* strain would be more resistant to cell wall-active antibiotics such as cephalosporins. Thus, we determined the minimum inhibitory concentration (MIC) of antibiotics whose tolerance is dependent upon CroR activation (e.g. vancomycin, ceftriaxone, and cefotaxime) in the *croR, croS* and Δ*srtA*Δ*htrA* mutant strains (**Fig 7D**) (37, 43, 50). In line with previous studies, the *croR::tn* strain was hyper-susceptible to ceftriaxone and cefotaxime, while the *croS::tn* strain was not (43). Surprisingly, the Δ*srtA*Δ*htrA* strain was highly sensitive to these cephalosporins (Δ*srtA*Δ*htrA* 16 and 4 µg/mL, WT>256 and >32 µg/mL) and inactivation of either *croR* or *croS* in this background further decreased antibiotic tolerance comparable to the single *croR*::tn strain. Vancomycin susceptibility was mainly observed in the context of *croR* inactivation.

Our results suggested that CroRS responds to endogenous, pilus-dependent membrane perturbation. We hypothesized that CroRS could also respond to exogenous membrane perturbations. The last resort antibiotic daptomycin perturbs the cell membrane, representing an exogenous source of membrane stress (51). RNA-Seq transcriptome analysis performed in our lab of WT OG1RF treated with subinhibitory concentrations of daptomycin induced expression of *htrA* and *croRS* (**Fig 7E, Table S7**), suggesting that CroRS might indeed also respond to exogenous membrane stress. Thus, to determine if CroRS contributes to daptomycin tolerance, we determined the daptomycin MIC of the panel of mutant strains. The single *croR::tn* and *croS::tn,* as well as the double Δ*srtA*Δ*htrA* strain were mildly more daptomycin sensitive. Strikingly, the triple *croS::tnΔsrtAΔhtrA* strain displayed a daptomycin hyper-susceptibility (*croS::tnΔsrtAΔhtrA* 0.032 µg/mL, WT 0.75 µg/mL) that was not apparent in the triple *croR::tnΔsrtAΔhtrA* (0.25 µg/mL). Overall, our results indicate that dual inactivation of *srtA* and *htrA* reduces the cephalosporin and daptomycin tolerance of *E. faecalis*, and that inactivation of the *croS* histidine kinase gene in this background further increases daptomycin susceptibility.

## Discussion

During colonization and infection, bacteria face unfavorable environmental conditions such as acidic pH and bile salts that cause proteins to misfold and aggregate, leading to substantial proteotoxic stress if left unresolved (13). This is especially true for surface exposed proteins that directly interact with the host milieu. To avoid accumulation of defective proteins that might impair crucial cellular processes, bacteria evolved protein quality control systems that clear aberrant proteins from the cell surface (21). To better understand how enterococci adapt to proteotoxic cell envelope stress, we used the *E. faecalis* Ebp pilus system. Here, we show that the HtrA chaperone acts as a protein quality control factor involved in clearance of aberrant membrane-anchored pili. By ensuring that pili do not accumulate on the membrane, HtrA acts as a first line of defense that prevents perturbation of membrane homeostasis and unproductive activation of CroRS, which promotes continued pili expression (i.e. increased proteotoxic stress) and profound cell morphology alterations.

The HtrA protease family plays a key role in protein quality control, including degradation of misfolded and mislocalized pili (21). Here we propose that in *E. faecalis*, HtrA monitors pilus biogenesis and sorting by SrtA and only acts in the presence of protein defects. This is exemplified by the fact that absence of *htrA* alone does not affect pilus biogenesis under optimal laboratory growth conditions. Our results using a protease-defective HtrA demonstrate that clearance of aberrant membrane-anchored Ebp is carried out by the *E. faecalis* HtrA chaperone rather than by direct proteolytic degradation. The exact molecular mechanism of HtrA-mediated pilus clearance is currently unknown, but it might involve interaction with other proteases or release of pili to the extracellular environment.

Many essential processes occur at the membrane, ranging from nutrient exchange and energy production, to interaction with the environment and protection from external insults. Thus, membrane homeostasis needs to be maintained to support proper functioning and cell viability. Aberrant accumulation of off-pathway pili triggers a membrane stress response in *E. coli* (15). Here we show that accumulation of membrane-bound pili also elicits a membrane stress response in *E. faecalis* that partially parallels the response in *E. coli* (32). Specifically, membrane overloading with pili in *E. faecalis* repressed non-essential lipoproteins (e.g. PTS and ABC-type transporters) and activated the aggregate-resolving ClpB chaperone as well as peptidoglycan-targeting enzymes (e.g. penicillin-binding proteins, peptidoglycan-active cell division genes, murein hydrolase regulators). By contrast, *E. faecalis* did not upregulate genes involved in translation. It is possible that in Gram-positive bacteria such as *E. faecalis*, accumulation of membrane-bound pili elicits a milder membrane stress response than in Gram-negative bacteria. Here, it appears that the cell tries to return to membrane homeostasis by decreasing the non-essential protein load of the membrane, while redirecting cell resources into modifying the cell wall. It is possible that parallel upregulation of *liaXYZ* (but not *liaFSR*) prepares the cell to activate the LiaFSR membrane stress response if required. Notably, bacterial stress responses are known to hierarchically orchestrate adaptations depending on the degree of stress perceived, and the inability to activate CroRS was shown to constitutively induce *liaX* transcription (36). It is interesting to note that the initial proteotoxic stress in our study (i.e. inability to sort pili to the cell wall due to inactivation of *srtA*) transitioned into a membrane stress response due to the inability to remove membrane-bound pili by the HtrA quality control system. Indeed, deletion of *srtA* or *htrA* elicited only few transcriptional changes. Altogether, these results highlight the importance of SrtA and HtrA to avoid escalating a localized proteotoxic stress to a more generalized membrane stress.

An intriguing finding of this work is that the Δ*srtA* and Δ*srtA*Δ*htrA* strains produced more pili than the WT strain, and that this trait was dependent upon CroR activation. This is counterintuitive at first, since proteotoxic stress responses typically remove mistargeted proteins by upregulating chaperones and proteases, while downregulating synthesis of non-essential proteins (e.g. Ebp pilus) (13). If CroRS evolved in part to counteract pilus-dependent membrane stress similar to *E. coli* Cpx, it would be expected to promote repression of *ebp* and activation of chaperones such as *htrA*. However, our data shows that CroR activation due to membrane-bound pili drives pilus expression, consequently perpetuating the source of endogenous stress. It is likely that CroRS is not directly involved in the response to proteotoxic stress. However, by inactivating the HtrA quality control system that normally clears proteotoxic stress, we uncovered a mild membrane stress response that is partially regulated by the CroRS TCS. In this scenario, CroR-dependent induction of pili and downstream biofilm formation could represent a protection strategy against membrane stressors. Overall, this leads us to propose the following model. During colonization and infection, pili are highly expressed but can be misfolded and aggregate as a result of harsh host conditions, impairing proper processing by sortase A (2, 7, 8, 13). Aberrant pili are continuously monitored and cleared (most likely indirectly) by HtrA, which is sufficient to resolve any proteotoxic stress that might arise. If the stress, however, is left unresolved and escalates into membrane stress, it triggers activation of CroRS, leading to hyper-piliation and activation of peptidoglycan remodeling pathways as protection mechanisms. It is currently unknown how CroS might sense cell membrane perturbations. In the closely-related *Streptococcus gordonii*, the TCS SGO_1180 was shown to monitor sortase A-dependent adhesin processing by directly sensing the remnant C-terminal LPXTG sorting motif that remains inserted in the membrane after efficient cell-wall anchoring (52). Here, accumulation of the C-peptide that remains after sortase A processing inhibits activation of the TCS HK. BLASTP analysis suggests that SGO_1180 is not a CroS ortholog. However, it will be interesting to see if lack of Ebp C-pep is directly or indirectly linked to partial mobilization of the CroR regulon.

The crucial role that HtrA plays in preventing unproductive CroR activation due to membrane-bound pili is exemplified by the absence of a stress response in the hyper-piliated Δ*srtA* strain. Aberrant activation of cell envelope stress responses are often associated with fitness costs for the cell (15, 42), and CroR phosphorylation specifically has been shown to require tight control to avoid deleterious fitness defects (43), which argues in favor of not activating CroR unless required. This is especially true for *E. faecalis* OG1RF, a strain that harbors the CisRS TCS capable of phosphorylating CroR (43). Therefore, it is possible that *E. faecalis* makes use of the HtrA quality control system to efficiently clear toxic pilus aggregates without eliciting a membrane stress response. While our *in vivo* results suggest that HtrA promotes persistent wound colonization of *E. faecalis*, further studies are required to evaluate the significance of HtrA and CroRS to enterococcal colonization and infection.

The CroRS TCS of *E. faecalis* has been predominantly studied in the context of cell wall-damaging antibiotics, where CroRS is a key determinant in *E. faecalis* tolerance to cephalosporins (e.g. ceftriaxone). Recently, CroRS was shown to be important for cell wall homeostasis in the absence of antibiotics (36). Our proposed model broadens the scope of the CroRS TCS from a cell wall stress responder to also a membrane stress responder. Basal cephalosporin tolerance in *E. faecalis* is primarily mediated by the CroR-dependent Pbp4 and PbpA (35). Considering that CroR is active, and *pbp4* and *pbpA* transcription is induced in the Δ*srtA*Δ*htrA* mutant, it was puzzling that the strain was hyper-sensitive to cephalosporins nearing *croR::tn* values. However, the observation that inactivation of *croS* in this background did not revert the phenotype argues that the Δ*srtA*Δ*htrA* strain cannot mount a full CroR response on the cell surface. It is possible that either absence of *srtA* or *htrA*, or that membrane perturbation in the double mutant strain impairs proper localization or enzymatic function of membrane-bound penicillin binding proteins such as Pbp4 and PbpA, rendering the cell less tolerant even with an active CroR regulon and an induced *pbp4* and *pbpA* transcription. Interference with penicillin-binding proteins could also explain the slight daptomycin susceptibility of the Δ*srtA*Δ*htrA* strain. Notably, work in *Staphylococcus aureus* showed that alterations in Pbp2 membrane localization determines susceptibility to β-lactams in constitutive daptomycin-resistant (i.e. LiaR active) cells, indicative of an overlap between CroRS and LiaFSR at the level of penicillin-binding proteins (53). In this regard, the hyper-sensitivity of the triple OG1RF *croS::tnΔsrtAΔhtrA* is reminiscent of LiaFSR defective mutants (42).

In this study we have uncovered an intricate link between the HtrA chaperone, the Ebp virulence factor, and the antibiotic response system CroRS (**Fig. 8**). Our results confirm the role of HtrA as a pilus quality control system in *E. faecalis*, where it contributes to clearance of membrane-bound Ebp pili and avoids downstream membrane stress. We show that the CroRS TCS can be activated by endogenous (off-pathway pili) and possibly also by exogenous (daptomycin) membrane stress (as opposed to classic antibiotic-mediated cell wall damage), and that this leads to RodA-dependent cell morphology alterations and to pilus expression in an EbpR-independent manner. Further studies will be required to shed light on the role of HtrA during *E. faecalis* proteotoxic stress during colonization and/or infection and to elucidate its connection to CroRS.

**Fig 8.**
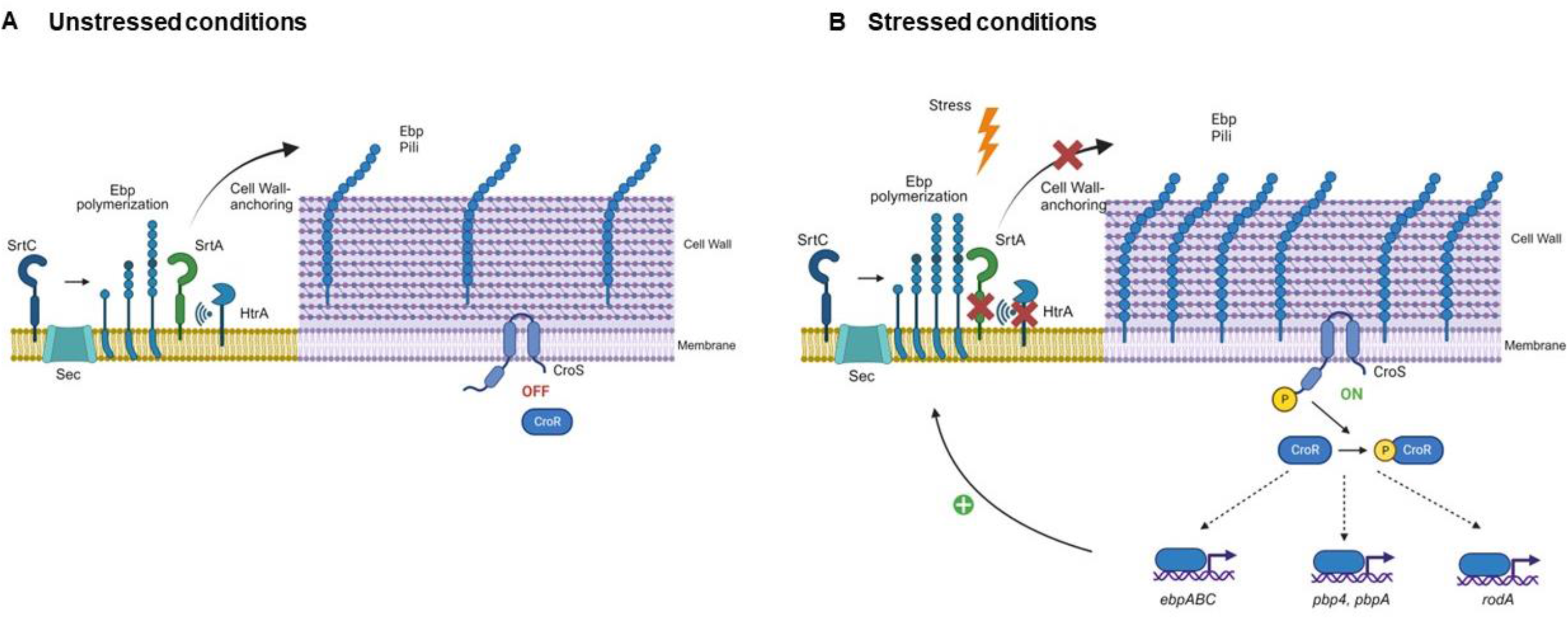
Model integrating the HtrA quality control system and the CroRS signalling pathway into pilus biogenesis. **(A)** During unstressed conditions, pilus monomers are exported through the Sec pathway, polymerized on the cell membrane by sortase C (SrtC), and efficiently anchored to the cell wall by sortase A (SrtA) (27). The HtrA bifunctional chaperone/protease monitors pilus processing by SrtA and contributes to clearance of sporadic aberrant pili. **(B)** Certain environmental stresses (e.g. stomach pH, gut bile salts, immune oxidative burst) cause protein misfolding and aggregation, impairing successful processing by SrtA. If HtrA activation is insufficient to cope with the proteotoxic stress, aberrant proteins accumulate and perturb the cell membrane, leading to activation of the CroS histidine kinase. CroS in turn activates the CroR response regulator, which regulates transcription of pilus (*ebpABC*), penicillin-binding proteins (*pbp4, pbpA*), and cell division (*rodA*) genes among others. Dotted arrows indicate confirmed indirect regulation (*pbpA*) (64) or unknown direct or indirect regulation by CroR. Induction of *ebpABC* genes leads to hyper-piliation, while repression of *rodA* partially leads to cell morphology alterations.

## Materials and methods

### Bacterial strains and general growth conditions

Bacterial strains and plasmids used in this study are listed in **S1 Table**. We inoculated *E. faecalis* from single colonies and grew cells statically at 37°C in Brain Heart Infusion (BHI) broth (Acumedia, USA) or agar (1.5%, Difco, US) for all assays unless otherwise stated. *E. coli* was grown in Luria Bertani (LB) broth (Difco, US) with shaking, or on agar plates at 37°C for DNA isolation and manipulation. To stimulate pilus expression in *E. faecalis,* strains were grown statically in Trypticase Soy Broth (Oxoid, UK) supplemented with 0.25% glucose (TSBG) and incubated at 37°C (2). We used Müller Hinton (MH) broth and 1.5% agar to perform antibiotic susceptibility assays. All inoculations were cultured for 15 to 18 hours unless otherwise stated. When required, antibiotics were added at the following concentrations: for *E. coli,* Kanamycin (Km) 50 mg/L or Erythromycin (Em) 500 mg/L; for *E. faecalis* strains, Em 25 mg/L, Km 500 mg/L, Rifampicin (Rif) 25 mg/L, Chloramphenicol (Cm) 10 mg/L. All antibiotics were purchased from Sigma-Aldrich Corporation, USA.

### DNA manipulation and construction of deletion mutants and complement strains

We used the Wizard® Genomic DNA Purification Kit (Promega, USA) to isolate bacterial genomic DNA from *E. faecalis* and the PureLink® Quick Plasmid Miniprep Kit (Invitrogen, USA) to isolate plasmid DNA from *E. coli*. Primers used in the study are listed in **S2 Table**. Primer design was performed based on the annotated complete genome of *E. faecalis* OG1RF (NC_017316) (54). Amplification of all gene products was performed using Phusion High-Fidelity DNA Polymerase (Thermos Scientific, USA); screening and validation of DNA sequences were performed using *Taq* DNA Polymerase (New England Biolabs, USA). T4 DNA ligase and restriction enzymes were purchased from New England Biolabs, USA. Ligation and restriction digestion were performed per respective manufacturer’s protocol. Plasmids were created by infusion cloning, where the In-fusion enzyme (Clontech, USA) fuses PCR-generated sequences containing a 15 bp overlap with linearized vectors on both ends of the insert via homologous recombination.

In-frame deletion of *htrA* (OG1RF_12305, new locus tag OG1RF_RS11805) was created according to a previously described method (55). Briefly, we amplified regions approximately 450 bp upstream and downstream of the *htrA* gene from OG1RF using primer pair *htrA* sew-R/Δ*htrA* del-F for the upstream region and *htrA* sew-F/Δ*htrA* del-R for the downstream region. These products were sewed together and amplified using Δ*htrA* del-F/Δ*htrA* del-R. We cloned the 900 bp PCR amplicon into pGCP213, a temperature sensitive Gram-positive plasmid (56) at the PstI restriction site to generate deletion construct p*delta*-*htrA.* We transformed the deletion construct into wildtype, Δ*srtA*, *croR::tn,* and *croS::tn* by electroporation and transformants were selected at 30°C on BHI agar containing Em. Single colonies were inoculated into individual tubes of BHI broth containing Em and passaged for two days at 30°C. Chromosomal integrant of this temperature-sensitive plasmid were selected by passaging the culture at 42°C in BHI, in the presence of Em. Selection for excision of the integrated plasmid by homologous recombination was accomplished by growing the bacteria at 30°C in the absence of Em. Loss of *htrA* locus in Em sensitive bacteria was verified by PCR using Δ*htrA* del-F/Δ*htrA* del-R and *htrA* cp-F/Δ*htrA* del-R (screen). To create Δ*srtA*Δ*htrA, croS::tnΔhtrAΔsrtA,* and *croR::tnΔhtrAΔsrtA* we transformed p*delta-srtA* (57) into Δ*htrA*, *croS::tnΔhtrA* and *croR::tnΔhtrA,* respectively. Loss of *srtA* locus in Em sensitive bacteria was demonstrated by PCR using Δ*srtA* del-F/Δ*srtA* del-R. To obtain the quintuple Δ*srtA*Δ*ebpABCΔhtrA* strain, the deletion construct pdelta-*htrA* was transformed into the Δ*srtA*Δ*ebpABC* strain and screened for the absence of *htrA*. We constructed a *htrA* complementation vector by amplifying the *htrA* coding sequence plus 200 base pairs upstream of the *htrA* start codon to include its native promoter with *htrA* cp-F/*htrA* cp-R or *htrA* cp-F/*htrA* cp-R-HA. We digested the resultant 1.5 kb PCR product with EcoRI and BamHI and ligated into pGCP123, giving rise to p*htrA* or p*htrA-HA*. To construct a HtrA expression plasmid defective of protease activity, extracted p*htrA* was subjected to a single amino acid change on the conserved serine (S271) on *htrA* allele by site-directed mutagenesis (SDM) using S271A_F_SDM/S271A_R_SDM. To construct an EbpABC expression plasmid defective in pilus assembly, extracted p*ebpABCsrtC* was subjected to a single amino acid change on the conserved lysine (K186) of the *ebpC* allele using primer pairs EbpC_K186A_F/EbpC_K186A_R. All plasmids were sequenced for verification by standard Sanger sequencing (AIT biotech, Singapore) We verified the protein expression and stability of proteins in these expression plasmids by immunoblot using respective antibody immune sera (**S3 Table**).

### Growth kinetics at different temperatures

Overnight cultures were diluted 10-fold in BHI and grown for 1 h at 37°C. Cells were then normalized to OD_600_ of 0.003 in a final volume of 50 mL media. To test the growth kinetics under various temperatures, we incubated each set of cultures at 37°C or 42°C. Cell culture was extracted hourly for measurement of OD_600_ and colony forming unit (CFU/mL) plating on BHI plates. The plates were incubated at 37°C aerobically for 16 h.

### Environmental stress tolerance assays

We prepared BHI plates containing NaCl (0.5 to 1.5 M) or H_2_O_2_ (1.0 to 2.5 mM; Sigma-Aldrich). For growth experiments involving pH, we adjusted the initial pH of BHI agar to pH 5.5, 6.0, and 7.0 with HCl before sterilization. 50 mM citrate-phosphate buffer of the desired pH was added to media after sterilization. Overnight cultures were spun down and washed with sterile 1× PBS. Cultures were resuspended in either the same or 1/10 of the original volume in 1× PBS and measured at OD_600_ using a spectrophotometer (UVmini-1240, Shimadzu, Japan). Cells were normalized to OD_600_ of 0.5 and 10-fold serial dilutions were made. 5 µL aliquot of each dilution was spotted on BHI agar plates containing different stressors and plates were incubated at 37°C aerobically for 48-72 h.

### Biofilm Assays

Overnight cultures were normalized to OD_600_ of 0.5-0.6. Then, 8 µL of normalized culture were mixed with 200 µL of BHI or TSBG media in a 96-well plate (Nunc™ MicroWell™ 96-Well Microplates, Thermos Scientific™, USA). To test the effects of temperature on biofilm formation, the plates were incubated at 30°C, 37°C or 42°C statically for 24 or 48 h. Following incubation, planktonic bacteria were discarded by tipping content into a waste container. The plates were washed twice with 1 X PBS to remove non-adherent bacteria before staining with 0.1% crystal violet for 15 mins at 4°C. The plate was then washed with PBS until negative control is clear. 200 µL of ethanol: acetone (4:1) was added per well and incubated on a belly dancer for 30-60 mins with lid on to prevent evaporation. Absorbance was read at OD_595_ using the microplate reader (Infinite M200 Pro, Tecan, Switzerland). All biofilm assays were performed with three biological replicates, each with 12 technical replicates.

### Murine wound infection model

The murine wound infection model was carried out as described elsewhere (25). Groups of four to five male C57BL/6 mice (7-8 weeks old, 22 to 25 g; InVivos, Singapore) were used. Briefly, bacterial strains were grown in 15 mL BHI media for 16-18 h at 37°C. Cells were pelleted, resuspended in 5 mL sterile 1× PBS and the OD_600_ was normalized to 2 × 108 CFU/mL. For competitive infection, an equal volume of each strain was mixed prior to infection. Mice were euthanized at 8 or 72 hpi and one cm by one cm squared piece of skin surrounding the wound site was excised and collected in sterile 1× PBS. After homogenation, viable bacteria were enumerated by plating onto both BHI plates and antibiotic selection plates to ensure all recovered colony forming units corresponding to the inoculating strain. To measure the fitness of the strains in causing infection, we calculated the competitive index (CI) as described elsewhere (25).

### Bacterial cell fractionation

*E. faecalis* was grown to mid-log phase at the indicated temperatures and OD_600_ of 0.5, normalized to 0.6, and equivalent volumes subjected to centrifugation at 8,000 x *g* for 5 minutes. We washed the cell pellets once in PBS and digested for one hour with 10 mg/mL lysozyme from chicken egg white (Sigma-Aldrich, USA) in lysozyme buffer (10 mM Tris-HCL pH 8, 50 mM NaCl, 1 mM EDTA, 0.75 M Sucrose) yielding the whole cell lysate. For further fractionation, the lysate was further subjected to centrifugation at 20,000 x *g* for 5 minutes. The resulting supernatant containing material liberated from the cell wall digestion was designated the cell wall fraction and the pellet designated the protoplast fraction. All fractions were stored at -20°C until use.

### Immunoblotting

Whole cell lysates or bacterial fractions were boiled for at least 15 mins in NuPAGE® LDS Sample Buffer (4×) with dithiothreitol (DTT) and SDS-PAGE was performed with NuPAGE Novex 3 to 8% Tris-acetate gels in NuPAGE Tris-acetate SDS running buffer (Life Technologies Corp., USA) to resolve proteins >150kDa. For smaller proteins, NuPAGE Novex 4-12% Bis-Tris gels in NuPAGE MOPS SDS running buffer were used. The iBlot® Transfer Stacks were used to transfer proteins using the iBlot® Gel Transfer Device (Life technologies Corp. USA). We blocked the membrane with 3% P-Bovine Serum Albumin (P-BSA) for one hour at room temperature or at 4°C overnight and then incubated with the indicated anti-sera for two hours at room temperature or at 4°C overnight, with gentle shaking. Blots were washed and then incubated with Pierce^TM^ horseradish peroxidase-conjugated secondary antibodies (Thermo Fisher Scientific, Inc., USA) and incubated with Super Signal West Femto or Pico chemiluminescent substrate (Thermo Fisher Scientific, Inc., USA). We processed the Green X-ray film (Carestream, USA) with a Kodak processor (Kodak X-OMAT processor 2000). The PageRuler™ Prestained Protein Ladder, 10 to 180 kDa (Thermo Scientific, USA) was used to monitor protein sizes. Polyclonal antisera were generated commercially (SABio, Singapore) by immunization of hosts (**S3 Table**) with purified recombinant proteins, except for rabbit monoclonal anti-Hemagglutinin purchased from Thermos Scientific, USA. SecA, EbpA, and EbpB were generated previously (57).

### Bacterial cell preparation for immunofluorescence staining

*E. faecalis* strains grown in TSBG to mid-log phase were washed and normalized as described above. The cells were fixed with fresh 3% paraformaldehyde at 4°C for 10 mins and smeared on poly-L-lysine pre-coated slides (Polysciences, Inc., USA). Cells were washed once with 1× PBS and incubated with 100 times dilution of BODIPY® FL vancomycin (Van-FL) (Thermo Fisher Scientific, Inc., USA) at a final concentration of 5 ng/µL and incubated for one hour at room temperature in the dark. To visualize DNA, we added DAPI stain to the fixed cells at a final concentration of 2.5 ng/µL and incubated for 15 mins at room temperature in the dark. Cells were then blocked with filtered 2% P-BSA prior to adding 20 µL of respective primary antibody on to fixed cells and incubated overnight in 4°C, shaking. The primary antibody was paired with the respective Alexa Fluor® labelled secondary antibody and incubated at room temperature for one hour. Finally, we mounted the slide with mounting media (Vectashield®, USA) and coverslip for 30 mins before imaging or stored at 4°C in the dark prior to imaging with super-resolution structured illumination microscopy (SR-SIM) (Carl Zeiss, Germany) or inverted epi-fluorescence microscopy (Zeiss Axio Observer Z1, Germany). To visualize chain length, 5 µL of the culture was mixed with an equal volume of low melting agar (BioWorld, USA) on a glass slide before covering it with a coverslip. The slides were visualized using a phase contrast microscope (Zeiss Axio Observer Z1; Carl Zeiss GmbH) fitted with a 100× oil immersion objective with a numerical aperture 1.4 optovar 1.0 magnification changer 1.5×. Images were collected with AxioVision (Carl Zeiss Zen 8.0 and analyzed with ImageJ (http://rsb.info.nih.gov/ij/).

### Transmission electron microscopy

For ultrastructural analysis, we grew *E. faecalis* strains overnight and sub-cultured 1:10 into 20 mL TSBG media and grew cells to mid-log phase. We fixed the bacteria in 2% paraformaldehyde/ 2.5% glutaraldehyde in 100 mM phosphate buffer, pH 7.4 for one hour at room temperature, washed in phosphate buffer, and post-fixed in 1% osmium tetroxide (Polysciences Inc.) for one hour. Samples were then rinsed extensively in deionized water prior to bloc staining with 1% aqueous uranyl acetate (Ted Pella Inc., Redding, CA) for one hour. Following several rinses in deionized water, we dehydrated the samples in a graded series of ethanol and embedded in Eponate 12 resin (Ted Pella Inc.). Sections of 95 nm were cut with a Leica Ultracut UCT ultramicrotome (Leica Microsystems Inc., Bannockburn, IL), stained with uranyl acetate and lead citrate, and viewed on a JEOL 1200 EX transmission electron microscope (JEOL USA Inc., Peabody, MA) equipped with an AMT 8-megapixel digital camera and AMT Image Capture Engine V602 software (Advanced Microscopy Techniques, Woburn, MA).

### RNA purification for RNA sequencing

For RNA-Seq analysis of daptomycin-treated OG1RF WT, sample preparation was adapted from a previous study (58). Briefly, bacterial overnights were sub-cultured in BHI + 50 mg/L CaCl_2_ and incubated to OD_600_ ∼0.4 (log phase). Then, cells were exposed to sub-inhibitory concentrations of daptomycin (1µg/mL) for 15 min prior to RNA extraction. RNA extracted from untreated cells served as a control. For mRNA transcriptomic analyses of OG1RF WT, Δ*htrA*, Δ*srtA*, Δ*srtA*Δ*htrA* and *croR*::TnΔ*srtA*Δ*htrA* we grew the bacteria overnight, statically in TSBG media at 37°C. The next morning, cultures were diluted 1:10 and grown to an optical density (600 nm) of 0.5. Total RNA was extracted using the UltraClean® Microbial RNA Isolation Kit (MO BIO Laboratories Inc., Singapore). Extracted RNA samples were subjected to rigorous DNase treatment using TURBO DNA-free™ kit (Ambion®, Singapore) and purified DNA-free RNA samples were subjected to ribosomal depletion with Ribo-Zero™ Magnetic Kits (Epicentre®, Singapore), all according to manufacturer’s protocols. Quantification of RNA and DNA were performed using Qubit™ RNA Assay Kits and Qubit™ dsDNA HS Assay Kits (Invitrogen, Singapore), respectively. The integrity of RNA was analyzed by gel electrophoresis using Agilent RNA ScreenTape (Agilent Technologies, Singapore). RNA samples were prepared in triplicate from three independent biological samples. mRNA libraries for RNA sequencing were prepared using TruSeq Stranded mRNA Library Prep Kit (Illumina, USA), the quality of the library analyzed via Bioanalyzer (Agilent, USA), and sequencing performed using an Illumina Miseq V2 machine. RNA sequencing reads were mapped to the *E. faecalis* OG1RF reference genome (NCBI accession: NC_017316.1) using BWA (v0.5.9) with default parameters (59, 60). Sequencing reads (Accession numbers CP002621.1) mapping to predicted open reading frames (ORFs) were quantified using HTSeq (61). Counts for ribosomal and transfer RNA sequences were filtered out of the data set and differential expression analyses were performed in R (version 2.15.1) using the Bioconductor package, edgeR (62). Significantly differentially expressed genes were determined using a P-value and false discovery rate (FDR) cut-off of 0.05. We annotated differentially expressed genes using a combination of KEGG annotations, as well as manual annotation using operon and other functional data from the literature and *E. faecalis* OG1RF database available on BioCyc https://biocyc.org/<u>) into</u>.

### Quantitative Real-time Polymerase Chain Reaction (qRT-PCR)

Quantitative reverse transcriptase PCR (qRT–PCR) was performed using a two-step method. 4900 ng of DNA-depleted RNA was first converted to complementary DNA (cDNA) using Superscript® III First-Strand Synthesis System Kit (Invitrogen, USA). Following cDNA synthesis, 0.09 ng of cDNA per well was used in qRT-PCR with KAPA SYBR® FAST qPCR Master Mix Kit (2×) (KAPA Biosystems, USA) on an Applied StepOnePlus^TM^ Real-Time PCR System (Applied Biosystems, USA). No amplification was observed for no-template control in qPCR reaction (C_T_ value above 35). To compare the differences between the target genes, the ΔΔC_T_ method was used (Livak & Schmittgen, 2001). Prior to the ΔΔC_T_ analysis, qPCR data was validated by running a standard curve for each gene as described in Applied Biosystems User Bulletin No.2 (P/N 4303859) and elsewhere (Livak & Schmittgen, 2001). The housekeeping gene gyrase B (*gyrB*) was used as an endogenous control in this study (Djoric & Kristich, 2015, 2017). Melting curve analyses were employed to verify the specific single-product amplification. Primers used in the study are listed in **S2 Table** and were generated using NCBI primer design software (Primer-BLAST) to amplify PCR products of size between 100-150 bp.

### Antibiotic susceptibility determinations

E-test (ETEST®, BioMérieux, USA) MICs were determined with 1.5% MH agar using the incubation condition as described above. Briefly, bacteria from stationary phase cultures in MHB media were washed and normalized visually in 1× PBS to McFarland standard 1.0. E-test inoculum preparation, plating, strip application, and MIC determinations were carried out according to the manufacturer’s protocol (63).The antibiotics tested were vancomycin, daptomycin, ceftriaxone, and cefotaxime. Insufficient growth of bacteria on the agar plate to form a bacteria lawn for accurate MIC determination after 18 h incubation will be given an additional incubation time of not more than 24 h.

### Statistical analyses

Data from multiple experiments were pooled. Statistical significance for biofilm assay was determined using a two-tailed unpaired t-test. Statistical significance for in vivo animal experiments was determined using one-way ANOVA, Kruskal–Wallis test with Dunn’s post-test to correct for multiple comparisons. Statistical significance with the relative density of protein level was determined using Tukey’s multiple comparisons test. Statistical significance with the percentage of cells expressing pilin was determined using the Holm-Sidak method. Statistical significance with the number of division septa per cell unit was determined using the Holm-Sidak method. Statistical significance with the qRT-PCR was determined using Tukey’s multiple comparisons test. Unless otherwise stated, values represented means ± SEM derived from at least three independent experiments and/ or three technical replicates. * P ≤ 0.05; ** P ≤ 0.01; *** P ≤ 0.001; **** P ≤ 0.0001; P ≥ 0.05, differences not significant (ns). GraphPad Prism 7 software (GraphPad Software, La Jolla, CA) was used for statistical analyses.

### Ethics Statement

We performed all approved procedures in accordance with the Institutional Animal Care and Use Committee (IACUC) in Nanyang Technological University, School of Biological Sciences (ARFSBS/NIEA0198Z) for murine wound infection model.

### Data availability

All the sequencing datasets have been deposited in the National Center for Biotechnology Information (NCBI) Gene Expression Omnibus (GEO) database under accession number (*accession number pending*).

## Acknowledgements

We thank Gary Dunny and Jennifer Dale for providing the *E. faecalis* transposon mutants used in this study, and Christopher Kristich for providing the Δ*cisS*Δ*croS* strain.

